# Efficacy of photodynamic therapy using 5-aminolevulinic acid-induced photosensitization is enhanced in pancreatic cancer cells with acquired drug resistance

**DOI:** 10.1101/2024.10.01.615935

**Authors:** Yiran Liu, Sally Kyei Mensah, Sergio Farias, Shakir Khan, Tayyaba Hasan, Jonathan P. Celli

## Abstract

The use of 5-aminolevulinic acid (ALA) as a precursor for protoporphyrin IX (PpIX) is an established photosensitization strategy for photodynamic therapy (PDT) and fluorescence guided surgery. Ongoing studies are focused on identifying approaches to enhance PpIX accumulation as well as to identify tumor sub-types associated with high PpIX accumulation. In this study, we investigated PpIX accumulation and PDT treatment response with respect to nodule size in 3D cultures of pancreatic cancer cells (Panc1) and a derivative subline (Panc1OR), which has acquired drug resistance and exhibits increased epithelial mesenchymal transition. In monolayer and 3D culture dose response studies the Panc1OR cells exhibit significantly higher cell killing at lower light doses than the drug naïve cells. Panc1OR also exhibits increased PpIX accumulation. Further analysis of cell killing efficiency per molecule of intracellular PpIX indicates that the drug resistant cells are intrinsically more responsive to PDT. Additional investigation using exogenous delivery of PpIX also shows higher cell killing in drug resistant cells, under conditions which achieve approximately the same intracellular PpIX. Overall these results are significant as they demonstrate that this example of drug-resistant cells associated with aggressive disease progression and poor clinical outcomes, show increased sensitivity to ALA-PDT.

## INTRODUCTION

PDT is a light-based treatment, in which wavelength-specific activation of a photosensitizing molecule leads to site-directed tumor destruction through photochemical processes (1). In PDT using 5-aminolevulinic acid (ALA) photosensitization, administration of exogenous ALA bypasses normal negative feedback regulation in heme biosynthesis, leading to accumulation of the photoactive heme precursor, protoporphyrin IX (PpIX) (1–5). Increased accumulation of PpIX in malignant relative to normal tissues has been associated with multiple factors, including lower activity of ferrochelatase and limited availability of iron in cancer cells (1).

In this study we specifically consider the role of drug resistance as a determinant of PpIX accumulation and PDT response. The acquisition of drug resistance is a common occurrence in disease progression, where initial responsiveness to chemotherapy gives way to refractory and recurrent disease as cancer cell populations with the ability to escape the cytotoxic effects of treatment become enriched by selection and adaptation (6, 7). Such survival mechanisms include increased expression of outward transport process transporters such as the well-known ABCG2 (8), and increased anti-apoptotic signaling. Cancer Stem Cells (CSCs) are also widely acknowledged for their role in drug resistance (9). Notably, PDT, under a range of conditions and in different disease settings has been shown to be able to bypass, and/or reverse drug resistance (9–14). PDT has also been shown to kill cervical cancer CSCs in vitro (9). Moreover, in the case of head and neck cancer, ALA-PDT has been found to improve the chemoresistance of CSCs (14). Additionally, studies have shown that drug-resistant pancreatic cancer cells are in fact more sensitive to photochemical treatment than their drug-sensitive counterparts (13).

In the present study, we examine PpIX accumulation and PDT response in Panc1 pancreatic ductal adenocarcinoma (PDAC) cells and a stable drug-resistant subline (Panc1OR) previously developed in our lab by exposure of Panc1 cells to increasing concentrations of oxaliplatin (11). This drug resistant sub-line has been shown to exhibit markedly increased epithelial-mesenchymal transition (EMT) and invasive behavior. Here we use PpIX quantification, viability assessment in monolayer and 3D cultures, and differential gene expression analysis from RNA sequencing data to identify determinants of ALA PDT responsiveness with respect to drug resistance status. In this context we particularly look at differential expression of ABCG2, genes relevant to heme biosynthesis, and genes related to oxidative stress response that could be protective against reactive oxygen species generated by PDT (15).

## MATERIALS AND METHODS

### Cell culture and reagents

Panc1 cells were obtained from ATCC. The stable drug-resistant line Panc1OR was generated by exposure to oxaliplatin chemotherapy as previously described (11). Cells were grown in T-75 cell culture flasks and incubated with appropriate media which also contained 10% fetal bovine serum (FBS), 100 IU/ml penicillin, and 0.5μg/ml Amphotericin B. All the cells were maintained in a humidified incubator at 37℃ with 95% air and 5% CO_2_.

### Monolayer cell culture

Cancer cells (3k∼5k/well) were grown in 96-well optically clear bottomed, black-walled plates (Ibidi; Fitchburg, WI) until reaching ∼90% confluence prior to treatment experiments.

### Matrigel based 3D cell culture

GFR (Growth factor reduced) Matrigel (Corning Life Sciences; Tewksbury, MA) was thawed at 4℃ before use. Matrigel beds were plated on culture dishes that had been stored in a -20℃ freezer overnight and maintained in contact with an icepack during plating. Liquid Matrigel was added to peri dish and the plate was gently shaken to create a flat surface while maintaining contact with the ice pack. The plate was then placed in the incubator at 37℃ for 30 minutes to allow gelation. Next, cancer cells were pipetted onto the Matrigel beds in each well and allowed to grow for 7 days to form 3D tumor nodules as previously described (16, 17). During this time, the 3D cell culture was incubated in media supplemented with 2% Matrigel.

### ALA solution

ALA powder (Thermo Fisher Scientific; Waltham, MA) was stored at -20℃. To prepare ALA solution, 0.0336g of powder was dissolved in 2ml of DPBS to obtain a 0.1M/L stock solution. The solution was then sterile filtered through a 0.22µm syringe driven filter unit and diluted to 2mM in the appropriate media prior to delivery to cell cultures.

### Photodynamic therapy treatment in vitro

PDT was performed using a low-cost portable fiber-coupled LED light source that was previously developed and reported by us (18–20). The specified experimental groups were incubated in media containing a 2mM ALA solution for a duration of 4 hours. Throughout this period, the cells were placed in an incubator and shielded with foil to protect from light. Just prior to treatment, the ALA media was substituted with regular media. The 635nm LED light was positioned vertically beneath the plate to activate the photosensitizer, with light dosages of 40J/cm^2^ at a fluence rate of 135mW/cm^2^. Following the treatment, the cells were once again shielded with foil and placed back into the incubator. Viability assessments were conducted 24 hours after treatment. In cases where photosensitization used direct exogenous delivery of PpIX the procedure was mostly the same except with incubation in the specified concentration of PpIX (Sigma-Aldrich, St. Louis, MO) solution for a duration of 4 hours. 24h later, viability is imaged and analyzed.

### Viability evaluation

24h after the treatment, the cells were stained with fluorescent vital dyes Calcein AM (Corning Life Sciences; Tewksbury, MA) and Ethidium Bromide. Stained cells were imaged at 5X magnification using a Zeiss AxioCam HRm camera mounted on a Zeiss AxioObserver.Z1 microscope (Carl Zeiss Microscopy GmbH, Jena, Germany). The live signal was based on Calcein AM fluorescence and dead signal was based on Ethidium Bromide fluorescence. Monolayer cell culture and 3D cell culture were quantified based on image segmentation using methods described previously (21).

### PpIX concentrations measurement

Cancer cells (200k/well) grown on 6-well plates for 5 days were photosensitized by incubation with ALA, or also KO143 (1µM, MedChemExpress; NJ) as described above prior to lysis in a chilled 0.25% Triton-X 100 solution (Thermo Fisher Scientific; Waltham, MA). Collected lysates were centrifuged at RCF=400g for 15 min to obtain a clear supernatant. The protein concentrations were quantified with the Pierce BCA assay (Thermo-Fisher Scientific; Waltham, MA). The PpIX fluorescence and protein absorbance were quantified with a Varioskan LUX Multimode Microplate Reader (Thermo Fisher Scientific; Waltham, MA). PpIX fluorescence is excited at wavelength of 405nm. Finally, the PpIX concentrations were calibrated using a standard curve of known PpIX concentration standards diluted in Triton X-100 solution.

## RESULTS

### ALA-PDT dose response in drug-resistant and drug-naïve PDAC cells

Investigation of dose dependent ALA-PDT response in cell cultures of Panc1 and their drug resistant counterparts, Panc1OR, exhibited a dramatic difference in response. Panc1OR, cells were significantly more sensitive to ALA-PDT than Panc1 all doses assessed, exhibiting almost complete cell killing at 40J/cm^2^ fluence (Figure 1).

**Figure 1.**
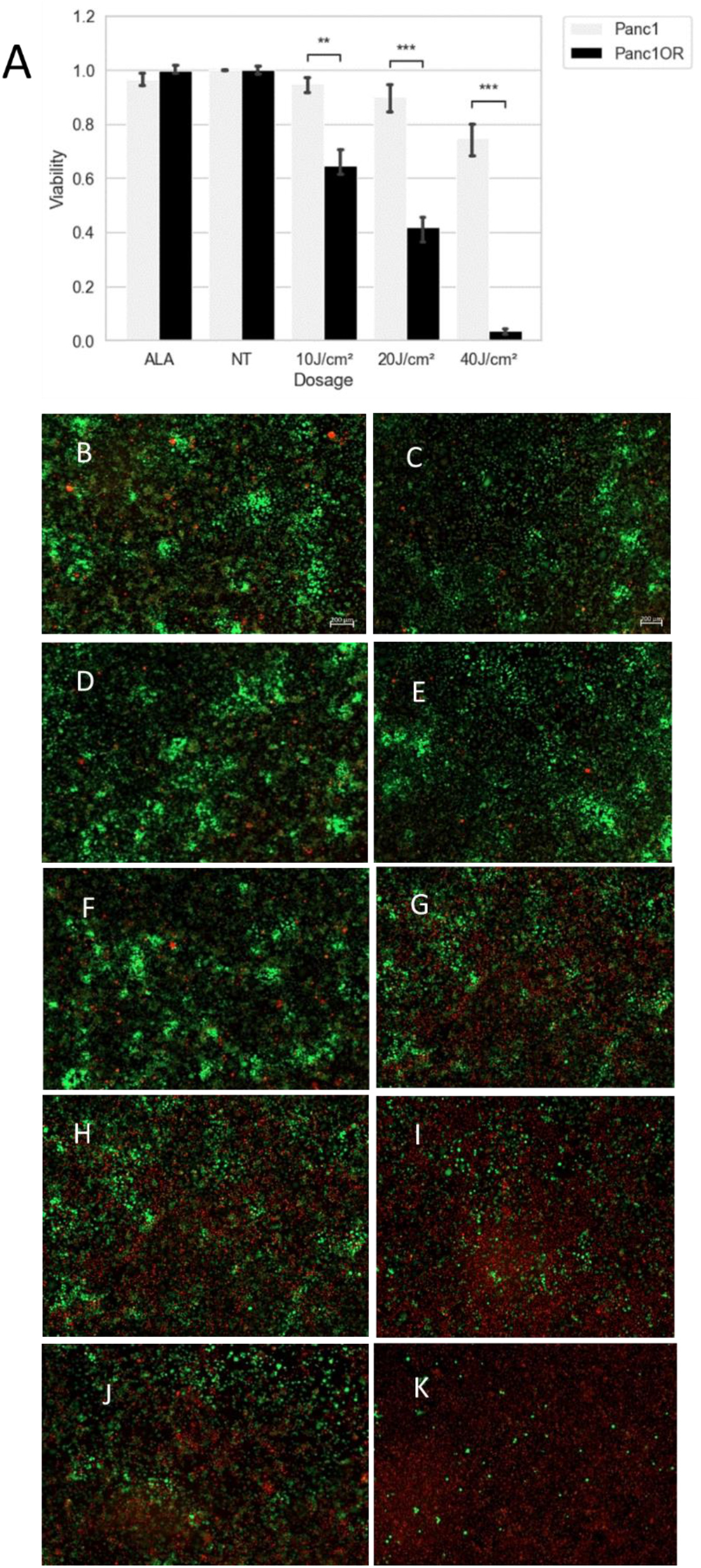
Cell culture stained with Calcein AM (green) and Ethidium bromide (red) after ALA PDT. A. ALA PDT response of Panc1 and Panc1OR monolayer cell culture. p-value annotation legend: ^*^: 1.00e-02 < p <= 5.00e-02. ^**^: 1.00e-03 < p <= 1.00e-02. ^***^: 1.00e-04 < p <= 1.00e-03. B. Panc1 monolayer cell culture had no treatment. D: Panc1 monolayer cell culture incubated with ALA but no treatment. F: Panc1 monolayer cell culture treated with 10J/cm^2^. H: Panc1 monolayer cell culture treated with 20J/cm^2^. J: Panc1 monolayer cell culture treated with 40J/cm^2^. C: Panc1OR monolayer cell culture had no treatment. E: Panc1OR monolayer cell culture incubated with ALA but no treatment. G: Panc1OR monolayer cell culture treated with 10J/cm^2^. I: Panc1OR monolayer cell culture treated with 20J/cm^2^. K: Panc1OR monolayer cell culture treated with 40J/cm^2^.

Matrigel based Panc1OR 3D cell cultures also exhibited higher sensitivity to ALA PDT when compared to Panc1 3D cell culture incubated under the same conditions (Figure 2). Given the same fluence (40J/cm^2^) after incubating with medium containing 2mM ALA for 4h, the viability of Panc1 3D cell culture is ∼5 fold higher than Panc1OR.

**Figure 2.**
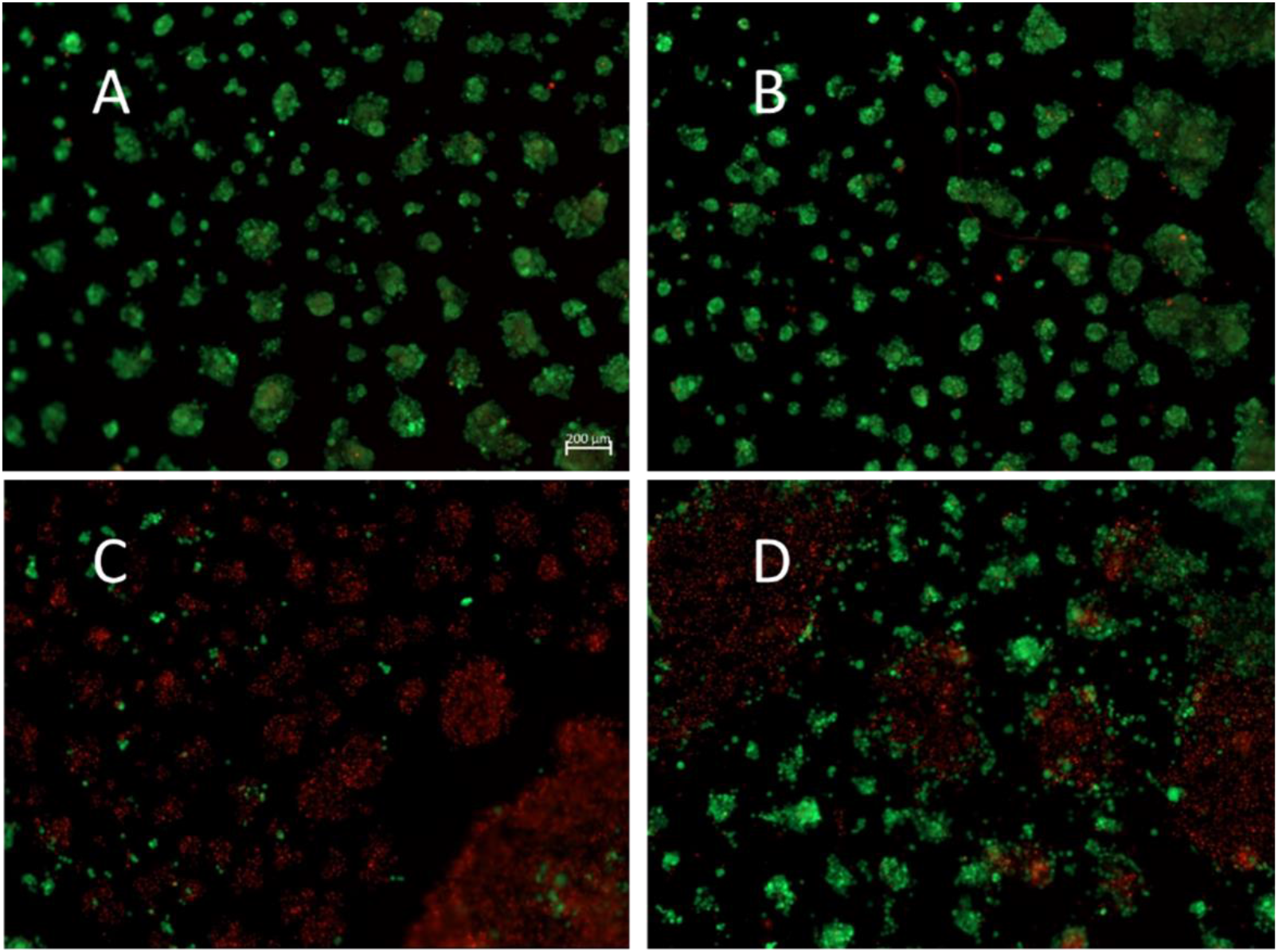
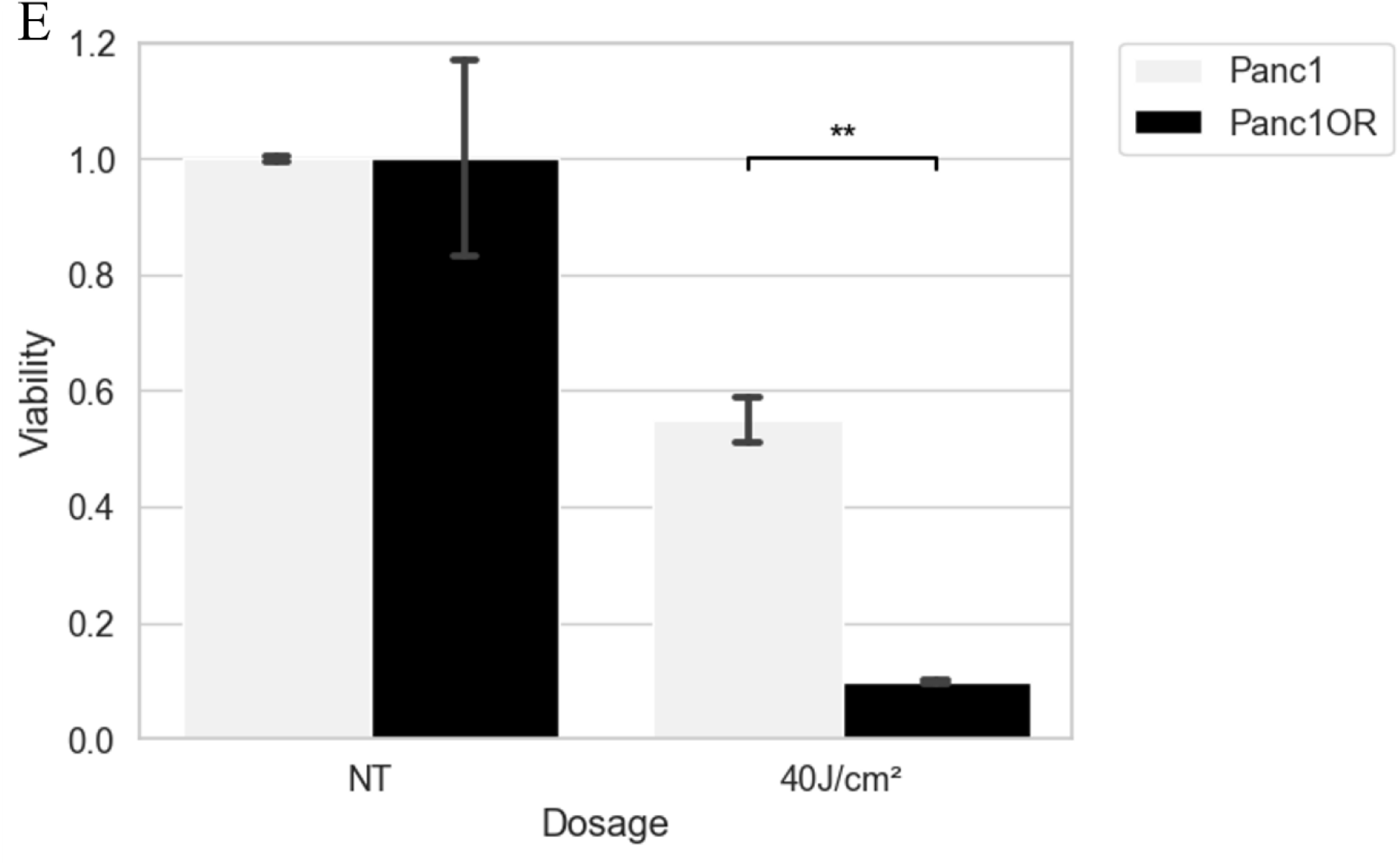
3D cell culture stained with Calcein AM (green) and ethidium bromide (red). A: Panc1OR 3D cell culture had no treatment. B: Panc1 3D cell culture had no treatment. C: Panc1OR 3D cell culture treated with 40J/cm^2^. D: Panc1 3D cell culture treated with 40J/cm^2^. E. ALA PDT response of Panc1 and Panc1OR 3D cell culture. p-value annotation legend: ^**^: 1.00e-03 < p <= 1.00e-02.

### Analysis of nodule-size dependent response to ALA-PDT in 3D PDAC cell cultures

For experiments using 3D cell cultures an imaging based methodology for analysis was used as described previously (21). The calcein and ethidium bromide fluorescence channels are both utilized to determine the nodule size, which are referred to as the live nodule size and dead nodule size, respectively. Areas below 10 pixels (2.39 µm^2^), were excluded from the analysis. The size of a nodule is determined by summing the live nodule size and dead nodule size. Viability is calculated by dividing the live nodule size by the sum of the live and dead nodule size.

There was no apparent distinction in the range of nodule sizes between Panc1 and Panc1OR (Figure 3.A, Figure 3.B). In the absence of treatment, the majority of nodules were found within the size range of a few to thousands µm^2^. As shown previously, treatment-induced breakdown of large nodules into smaller cell clusters was also noted in both Panc1 and Panc1OR (21). Without treatment, a significant number of live nodules are larger than 100 µm^2^. With treatment, the nodules larger than 100 µm^2^ diminishes substantially, but a dramatic increase in small nodules and single cells (size less than 100 µm^2^) is consistent with treatment-induced breakdown of large nodules into smaller ones.

**Figure 3.**
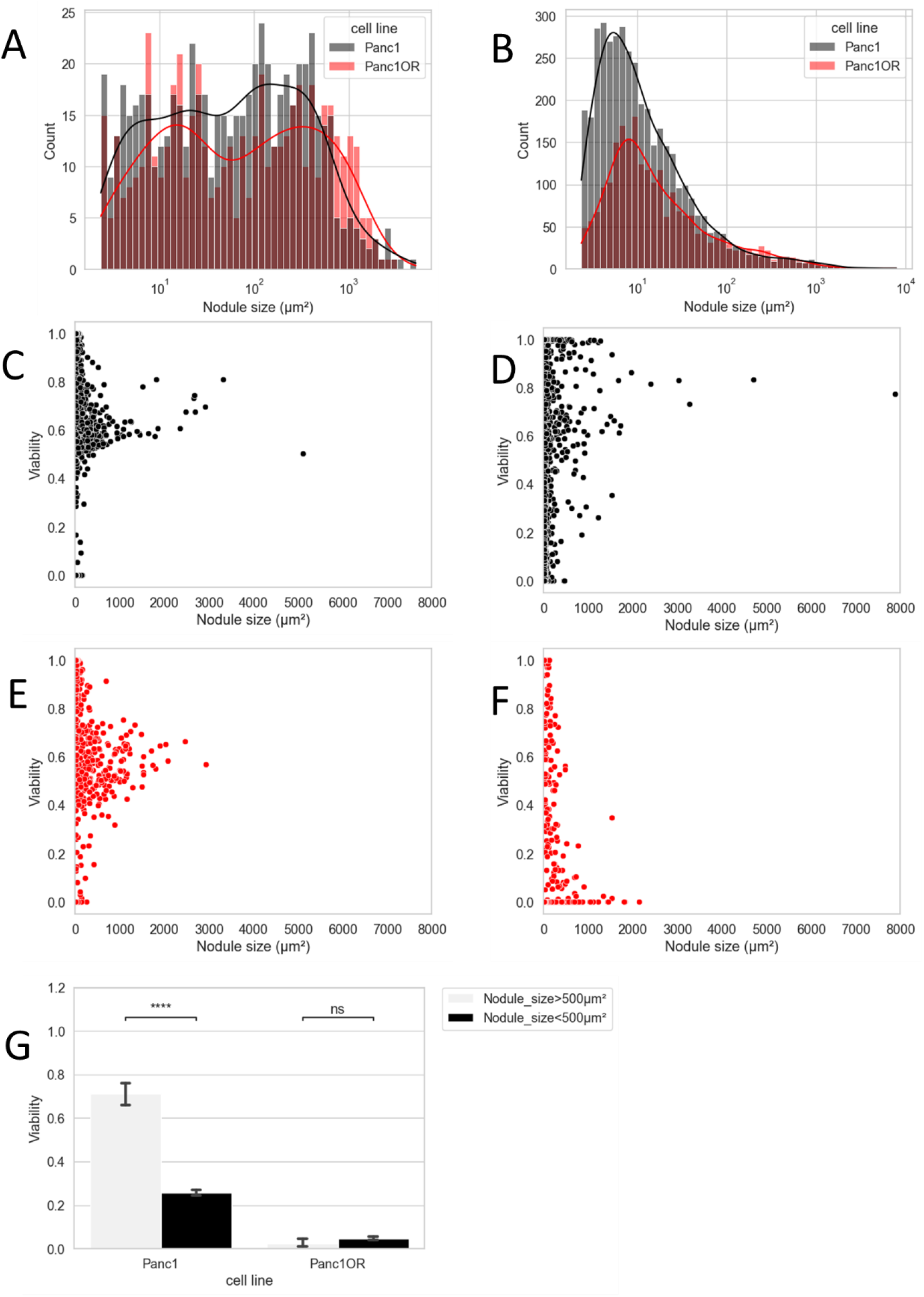
Heterogeneity of ALA PDT response in Panc1 and Panc1OR 3D cell culture. A. Nodule size range of Panc1 and Panc1OR 3D cell culture without treatment. B. Nodule size range of Panc1 and Panc1OR 3D cell culture with treatment. C. Viability - nodule size scatter of Panc1 without treatment. The size displayed on the horizontal axis and its associated viability on the vertical axis. D. Viability - nodule size scatter of Panc1 with treatment. It is assumed that small nodules (<500 µm^2^) showed higher responsiveness to ALA PDT compared to large nodules (>500 µm^2^), as large nodules associated with significantly higher viability than small nodules. E. Viability - nodule size scatter of Panc1OR without treatment. F. Viability - nodule size scatter of Panc1OR with treatment. The large nodules (>500 µm^2^) in Panc1OR had lower viability compared to small nodules (<500 µm^2^). This indicates that large nodules in Panc1OR seem to be more sensitive to the treatment. G. Comparison of viability associated with nodule size between Panc1 and Panc1OR. p-value annotation legend: ns: 5.00e-02 < p <= 1.00e+00. ^****^: p <= 1.00e-04.

The following scatter plots illustrating the correlation between the size of nodules and their viability. Each data point corresponds to a nodule, with its size displayed on the horizontal axis and its associated viability on the vertical axis. In the absence of treatment, the distribution of nodule sizes and associated viability in Panc1 (Figure 3.C) and Panc1OR (Figure 3.E) is remarkably similar to each other. However, with treatment (Figure 3.D, Figure 3.F), heterogeneity was observed in the relationships.

The response to treatment in Panc1 (Figure 3.D) has been found to vary depending on the size of the nodules. When compared to the untreated nodules, there was more presence of small nodules (<500 µm^2^) with lower viability. Additionally, large nodules (>500 µm^2^) associated with higher viability than small nodules. Consequently, it is assumed that for Panc1, smaller nodules show a higher responsiveness to ALA PDT compared to larger nodules.

Unlike Panc1, Panc1OR (Figure 3.F) displays a distinct pattern with larger nodules (>500 µm^2^) more likely to have lower viability in comparison to smaller nodules (<500 µm^2^). This suggests that larger Panc1OR nodules exhibit heightened sensitivity to the treatment. In Panc1, the viability of smaller nodules is lower than that of larger nodules, whereas in Panc1OR, this relationship is reversed, as illustrated in Figure 3.G.

### Quantification of PpIX accumulation

To explore the mechanism of increased ALA PDT response in Panc1OR, PpIX accumulation of Panc1 and Panc1OR monolayer cell culture was compared (Figure 4.A). The measurement confirmed the Panc1OR monolayer cell culture accumulated about twice the amount of PpIX of Panc1 monolayer cell culture when incubated in the medium contained 2mM ALA solution for 4h.

**Figure 4.**
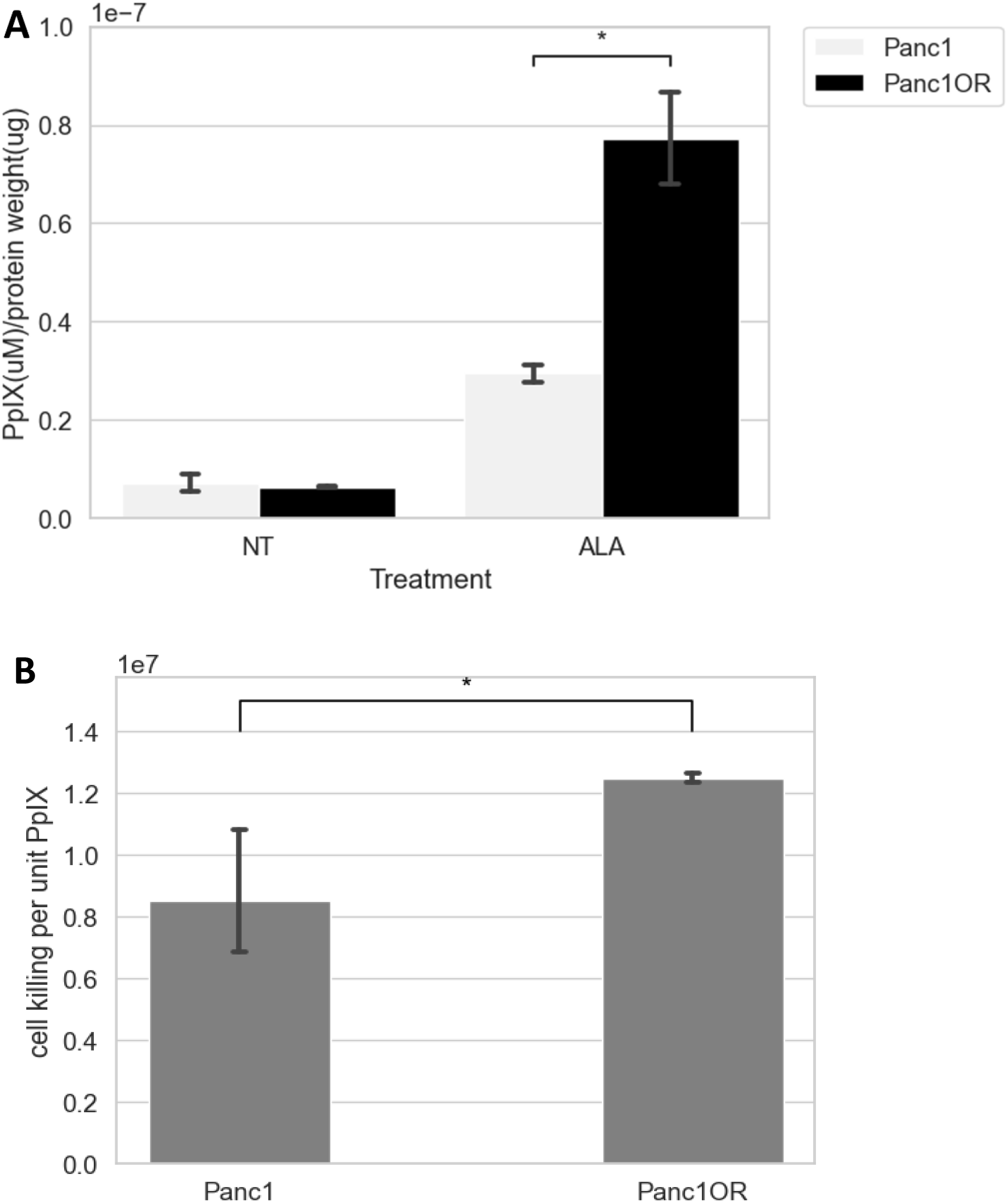
A: PpIX accumulation in Panc1 and Panc1OR monolayer cell culture. p-value annotation legend: ^*^: 1.00e-02 < p <= 5.00e-02. B: Comparison of cell killing per mole of PpIX between Panc1 and Panc1OR. p-value annotation legend: ^*^: 1.00e-02 < p <= 5.00e-02.

When examining the number of cell kill in monolayer cells which received 40J/cm^2^ treatment and PpIX concentrations following ALA incubation, Panc1OR exhibited a much higher number than Panc1 (Figure 4.B). Given that this correlation corresponds to the cell killing efficiency, it confirms that Panc1OR is more responsive to ALA PDT compared to Panc1.

### Response to PDT using exogenous PpIX photosensitization

To directly examine sensitivity to PpIX PDT independent of differences in heme biosynthesis, treatment response in Panc1 and Panc1OR monolayer cell culture photosensitized by exogenously delivered PpIX were compared. For these experiments cultures were incubated with PpIX and treated using the same 635nm LED light source. The results are compared and shown below (Figure 5). Given the same amount of PpIX concentrations and same fluence (20J/cm^2^), Panc1OR exhibits lower viability than Panc1, and this trend is consistent with varying the PpIX concentration.

**Figure 5.**
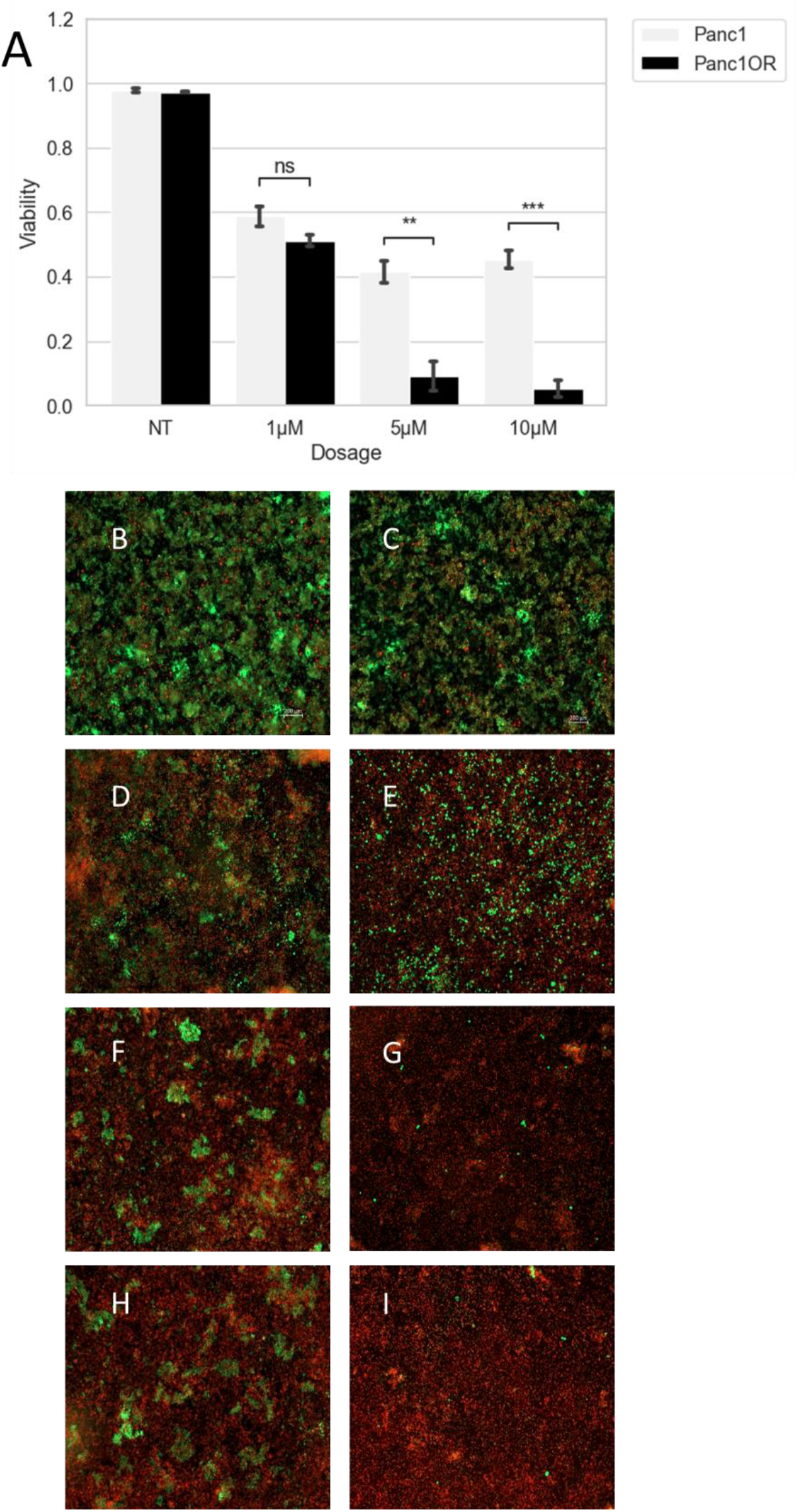
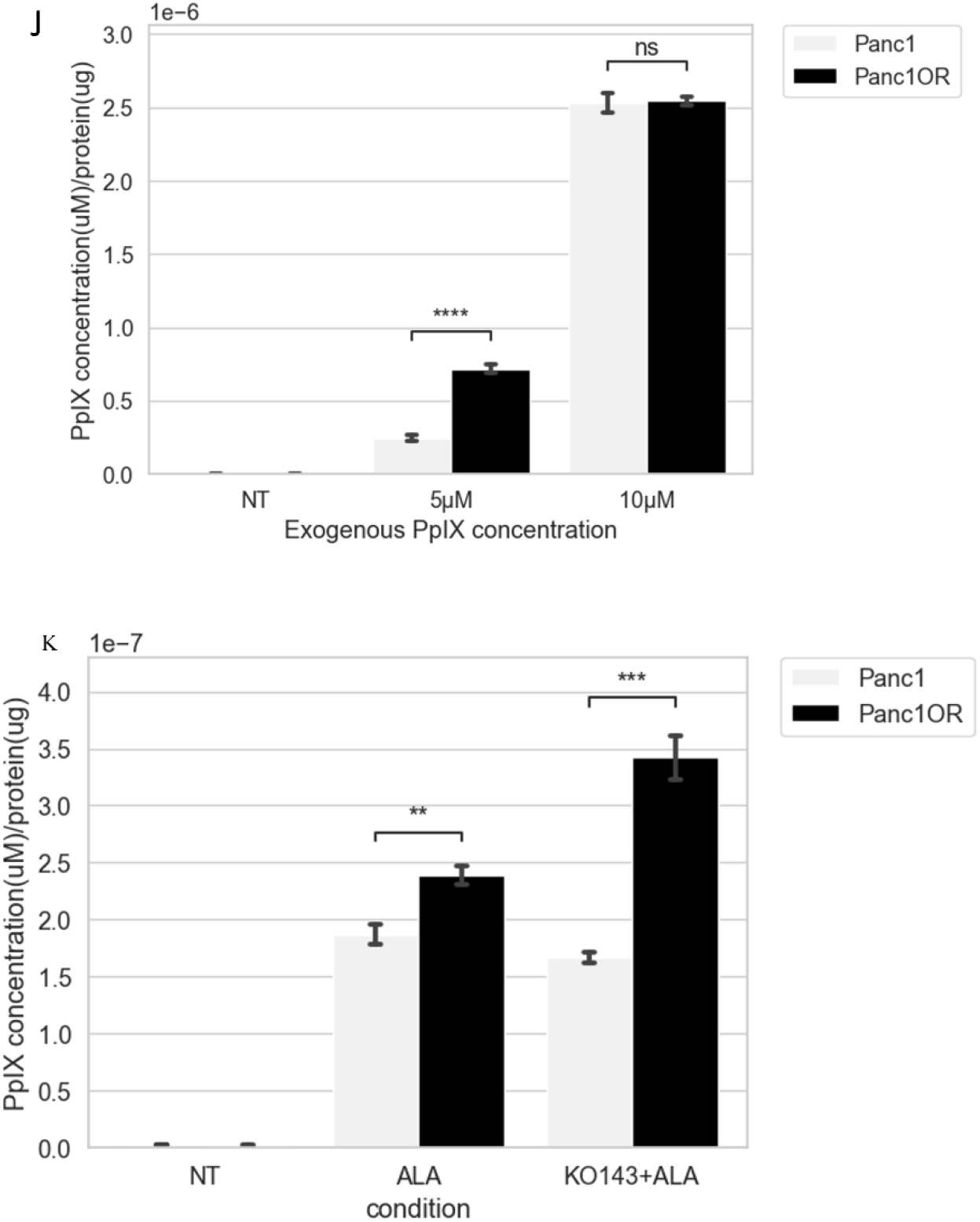
Panc1 monolayer cell culture and Panc1OR monolayer cell culture stained with Calcein AM (green) and Ethidium bromide (red) after PpIX PDT. A: PpIX PDT response of Panc1 and Panc1OR monolayer cell culture. p-value annotation legend: ns: 5.00e-02 < p <= 1.00e+00. ^**^: 1.00e-03 < p <= 1.00e-02. ^***^: 1.00e-04 < p <= 1.00e-03. B: Panc1 monolayer cell culture given regular medium. D: Panc1 monolayer cell culture given 1 µM PpIX and 20J/cm^2^ irradiation. F: Panc1 monolayer cell culture given 5 µM PpIX and 20J/cm^2^ irradiation. H: Panc1 monolayer cell culture given 10 µM PpIX and 20J/cm^2^ irradiation. C: Panc1OR monolayer cell culture given regular medium. E: Panc1OR monolayer cell culture given 1 µM PpIX and 20J/cm^2^ irradiation. G: Panc1OR monolayer cell culture given 5 µM PpIX and 20J/cm^2^ irradiation. I: Panc1OR monolayer cell culture given 10 µM PpIX and 20J/cm^2^ irradiation. J: PpIX accumulation of Panc1 and Panc1OR monolayer cell culture following exogenous PpIX incubation. ^****^: p <= 1.00e-04. K: PpIX accumulation of Panc1 and Panc1OR monolayer cell culture following ALA and ALA and ABCG2 inhibitor (KO143,1μM) incubation. p-value annotation legend: ^**^: 1.00e-03 < p <= 1.00e-02. ^***^: 1.00e-04 < p <= 1.00e-03.

Alternatively, we quantified the PpIX accumulation following the incubation of varied concentrations of exogenous PpIX. Given the significant difference in viability of Panc1 and Panc1OR under the 10μM PpIX incubation, it is unexpected to observe no variation in PpIX accumulation between Panc1 and Panc1OR when both are subjected to 10μM exogenous PpIX.

Combine these results all, it prompts the assumption that PpIX level alone may not be the sole factor contributing to the increased sensitivity to ALA PDT in Panc1OR, and suggests that Panc1OR are inherently more sensitive to PDT regardless of also having higher PpIX accumulation when photosensitized via ALA. This assumption is further supported by the fact that the enhancement persists even in the presence of the ABCG2 inhibitor (Figure 5.J). PpIX concentrations of Panc1OR and Panc1 are quantified following the incubation of ALA and ABCG2 inhibitor, and the results revealed that Panc1OR accumulated significantly more PpIX than Panc1 regardless of the variance in ABCG2 level.

## DISCUSSION

Here we show that drug-resistant PDAC cells exhibit a significantly higher sensitivity to ALA PDT compared to their drug-sensitive counterparts. Enhanced efficacy of PDT was observed in both monolayer and 3D cell cultures. We specifically discussed the cell killing per mole of PpIX following ALA PDT and ALA incubation, the results indicated that the number of Panc1OR is greater than Panc1 which confirmed the enhanced sensitivity in Panc1OR.

Comparison of the PpIX accumulation in Panc1 and Panc1OR indicated initially that the enhanced response could be partly due to increase in PpIX accumulation in the Panc1OR before the treatment. Since differential gene expression in Panc1OR relatively to the parental Panc1 was recently reported (36), we have the opportunity to further explore the underlying mechanisms driving these changes. While numerous highly significant DEGs were identified, only a small subset could be directly linked to heme biosynthesis and/or PpIX accumulation.

**Table 1.** Expression of genes involves PpIX formation and accumulation and transports in PANC1 and PANC1OR. The positive log2(FC) shows the genes that are upregulated in PANC1OR. The negative log2(FC) shows the genes that are downregulated in PANC1OR. Except ALAD, none of the other genes related to PpIX formation exhibits a significant difference between PANC1 and PANC1OR. PpIX transporter ABCG2 is upregulated in PANC1OR which is associated with enhanced efflux of PpIX and opposite to the observed higher PpIX accumulation.

| Symbol | Log2(FC) | p.value | Protein |
| --- | --- | --- | --- |
| UROS | -0.47103 | 0.220172 | Uroporphyrinogen-III synthase |
| UROD | 0.347245 | 0.571031 | Uroporphyrinogen decarboxylase |
| HMBS | -0.14441 | 0.846618 | Porphobilinogen deaminase |
| CPOX | 0.183069 | 0.754982 | Oxygen-dependent coproporphyrinogen-III oxidase |
| ALAD | 0.855181 | 0.042471 | Delta-aminolevulinic acid dehydratase |
| PPOX | -0.26084 | 0.718533 | Protoporphyrinogen oxidase |
| FECH | -0.39269 | 0.393977 | Ferrochelatase |
| ALAS1 | 0.130471 | 0.807286 | 5-aminolevulinate synthase, non-specific |
| SLC15A<br>2 | -0.62886 | 0.335944 | ALA uptake |
| ABCG2 | 2.916043 | 5.81E-21 | PpIX transporter |

Of the various genes directly involved in heme biosynthesis, only ALAD, which encodes delta-aminolevulinate dehydratase and catalyzes the second step of heme biosynthesis (22), shows a slight increase in expression in Panc1OR compared to Panc1, barely surpassing the p<0.05 cutoff. None of the other genes involved in heme biosynthesis exhibit a significant difference in expression between Panc1 and Panc1OR. (1, 23, 24)

Panc1OR exhibited a higher expression of ABCG2 compared to Panc1, which is not surprising for a drug resistant sub-line. ABCG2 functions as a PpIX transporter (26), and increased ABCG2 expression is typically associated with enhanced efflux of PpIX, leading to lower PpIX accumulation. However, in this case, the opposite trend is observed. While it has been demonstrated previously that PDT using verteporfin can decrease the expression of ABCG2 (30), we examined PpIX accumulation prior to light delivery, such that any PDT-dependent changes in ABCG2 would not yet have taken effect. Here we observed that even with the inhibition of ABCG2, Panc1OR cells exhibited higher PpIX levels compared to Panc1 cells following ALA incubation. Furthermore, in our experiments, the quantification of PpIX accumulation following exogenous PpIX incubation and viability assessment alongside PpIX PDT revealed that despite no variance in PpIX accumulation levels under certain conditions (10μM), the Panc1OR are still more responsive (Figure 5). It is also relevant here that PDT using mitochondrial-localizing photosensitizers is able to bypass anti-apoptotic signaling in cancer cells, as established by seminal works by Kessel and Olenick (31–33). Taken together our results suggest that increased responsiveness to PDT in the drug resistant cells is a combination of an intrinsic increase in sensitivity combined with increased accumulation of PpIX.

The drug resistant cells used in this study are very poorly differentiated, with fibroblast-like morphology and previously shown to express almost no E-cadherin at the protein level (11). It is noteworthy that this comports with recent clinical observations on differentiation status and PDT sensitivity. A recent study of PDT for oral squamous cell carcinoma (OSCC) in patients with early stage disease noted that more poorly differentiated lesions responded significantly better to PDT than well differentiated lesions (20).

Further investigation in this study looked at treatment response with respect to nodule size in our 3D cell cultures. ALA PDT treatment effectively fragmented the large nodules into smaller clusters in both Panc1 and Panc1OR. Given the observation of this phenomenon and the assumption that small nodules of Panc1 are more sensitive to ALA PDT, it suggests that pre-treatment with ALA-PDT could enhance response to subsequent chemotherapy or additional cycles of PDT as noted previously with verteporfin photosensitization in a similar 3D culture model (35).

Overall these results are significant in that drug-resistant cells, previously reported to exhibit marked increase in invasive behavior, are highly sensitive to PDT using ALA photosensitization. This fits in with a broader picture of the effectiveness of PDT against aggressive tumor cell populations, which are typically associated with poor clinical outcomes. The results presented in this report also contribute to the identification of specific phenotypic traits and tumor subtypes that are correlated with enhanced accumulation of PpIX, offering valuable insights for the development of innovative strategies to further enhance ALA-PDT in the future.

## ACKNOWLEDGMENTS

We would like to thank Dr. Gwendolyn Cramer, for originally establishing, characterizing and cryopreserving the drug resistant subline used in this study. This work was supported by funding from the National Cancer Institute, UH3 CA189901 (JPC and TH) and U01 CA279862 (JPC and TH). YL was supported by a College of Science and Mathematics Dean’s Doctoral Research Fellowship at the University of Massachusetts Boston.

## REFERENCES

1. Wachowska, M., Muchowicz, A., Firczuk, M., Gabrysiak, M., Winiarska, M., Wańczyk, M., Bojarczuk, K. and Golab, J. (2011) Aminolevulinic acid (ala) as a prodrug in photodynamic therapy of cancer. Molecules 16, 4140–4164. 10.3390/molecules16054140.

2. Petusseau, A. F., Bruza, P. and Pogue, B. W. (2022) Protoporphyrin IX delayed fluorescence imaging: a modality for hypoxia-based surgical guidance. J Biomed Opt 27. 10.1117/1.jbo.27.10.106005.

3. Boere, I. A., Robinson, D. J., de Bruijn, H. S., Kluin, J., Tilanus, H. W., Sterenborg, H. J. C. M. and de Bruin, R. W. F. (2006) Protoporphyrin IX fluorescence photobleaching and the response of rat Barrett’s esophagus following 5-aminolevulinic acid photodynamic therapy. Photochem Photobiol 82, 1638–44. 10.1562/2006-01-03-RA-763.

4. Kennedy, J. C., Marcus, S. L. and Pottier, R. H. (1996) Photodynamic therapy (PDT) and photodiagnosis (PD) using endogenous photosensitization induced by 5-aminolevulinic acid (ALA): mechanisms and clinical results. J Clin Laser Med Surg 14, 289–304. 10.1089/clm.1996.14.289.

5. Castano, A. P., Demidova, T. N. and Hamblin, M. R. (2004) Mechanisms in photodynamic therapy: Part one - Photosensitizers, photochemistry and cellular localization. Photodiagnosis Photodyn Ther 1, 279–293. 10.1016/S1572-1000(05)00007-4.

6. Aniogo, E. C., Plackal Adimuriyil George, B. [and Abrahamse, H. (2019) The role of photodynamic therapy on multidrug resistant breast cancer. Cancer Cell Int 19, 91. 10.1186/s12935-019-0815-0.

7. Vasan, N., Baselga, J. and Hyman, D. M. (2019) A view on drug resistance in cancer. Nature 575, 299–309. 10.1038/s41586-019-1730-1.

8. Chen, J., Mao, L., Liu, S., Liang, Y., Wang, S., Wang, Y., Zhao, Q., Zhang, X., Che, Y., Gao, L. and Liu, T. (2015) Effects of a novel porphyrin-based photosensitizer on sensitive and multidrug-resistant human gastric cancer cell lines. J Photochem Photobiol B 151, 186– 93. 10.1016/j.jphotobiol.2015.08.020.

9. Chizenga, E. P., Chandran, R. and Abrahamse, H. (2019) Photodynamic therapy of cervical cancer by eradication of cervical cancer cells and cervical cancer stem cells.

10. Duska, L. R., Hamblin, M. R., Miller, J. L. and Hasan, T. Combination Photoimmuno-therapy and Cisplatin: Effects on Human Ovarian Cancer Ex Vivo.

11. Cramer, G. M., Jones, D. P., El-Hamidi, H. and Celli, J. P. (2017) ECM Composition and Rheology Regulate Growth, Motility, and Response to Photodynamic Therapy in 3D Models of Pancreatic Ductal Adenocarcinoma. Mol Cancer Res 15, 15–25. 10.1158/1541-7786.MCR-16-0260.

12. Spring, B. Q., Rizvi, I., Xu, N. and Hasan, T. (2015) The role of photodynamic therapy in overcoming cancer drug resistance. Photochemical and Photobiological Sciences 14, 1476– 1491. 10.1039/c4pp00495g.

13. Lund, K., Olsen, C. E., Wong, J. J. W., Olsen, P. A., Solberg, N. T., Høgset, A., Krauss, S. and Selbo, P. K. (2017) 5-FU resistant EMT-like pancreatic cancer cells are hypersensitive to photochemical internalization of the novel endoglin-targeting immunotoxin CD105-saporin. Journal of Experimental and Clinical Cancer Research 36. 10.1186/s13046-017-0662-6.

14. Yu, C. H. and Yu, C. C. (2014) Photodynamic therapy with 5-Aminolevulinic acid (ALA) impairs tumor initiating and chemo-resistance property in head and neck cancer-derived cancer stem cells. PLoS One 9. 10.1371/journal.pone.0087129.

15. Pizzino, G., Irrera, N., Cucinotta, M., Pallio, G., Mannino, F., Arcoraci, V., Squadrito, F., Altavilla, D. and Bitto, A. (2017) Oxidative Stress: Harms and Benefits for Human Health. Oxid Med Cell Longev 2017. 10.1155/2017/8416763.

16. Celli, J. P., Rizvi, I., Evans, C. L., Abu-Yousif, A. O. and Hasan, T. (2010) Quantitative imaging reveals heterogeneous growth dynamics and treatment-dependent residual tumor distributions in a three-dimensional ovarian cancer model. J Biomed Opt 15, 1. 10.1117/1.3483903.

17. Evans, C. L., Rizvi, I., Celli, J., Abu-Yousif, A., de Boer, J. F. and Hasan, T. (2010) Abstract 3261: Visualizing treatment response dynamics of an in vitro three-dimensional ovarian cancer model. Cancer Res 70, 3261–3261. 10.1158/1538-7445.AM10-3261.

18. Hempstead, J., Jones, D. P., Ziouche, A., Cramer, G. M., Rizvi, I., Arnason, S., Hasan, T. and Celli, J. P. (2015) Low-cost photodynamic therapy devices for global health settings: Characterization of battery-powered LED performance and smartphone imaging in 3D tumor models. Sci Rep 5, 10093. 10.1038/srep10093.

19. Liu, H., Daly, L., Rudd, G., Khan, A. P., Mallidi, S., Liu, Y., Cuckov, F., Hasan, T. and Celli, J. P. (2019) Development and evaluation of a low-cost, portable, LED-based device for PDT treatment of early-stage oral cancer in resource-limited settings. Lasers Surg Med 51, 345–351. 10.1002/lsm.23019.

20. Siddiqui, S. A., Siddiqui, S., Hussain, M. A. B., Khan, S., Liu, H., Akhtar, K., Hasan, S. A., Ahmed, I., Mallidi, S., Khan, A. P., Cuckov, F., Hopper, C., Bown, S., Celli, J. P. and Hasan, T. (2022) Clinical evaluation of a mobile, low-cost system for fluorescence guided photodynamic therapy of early oral cancer in India. Photodiagnosis Photodyn Ther 38, 102843. 10.1016/j.pdpdt.2022.102843.

21. Celli, J. P., Rizvi, I., Blanden, A. R., Massodi, I., Glidden, M. D., Pogue, B. W. and Hasan, T. (2014) An imaging-based platform for high-content, quantitative evaluation of therapeutic response in 3D tumour models. Sci Rep 4, 3751. 10.1038/srep03751.

22. Ishida, N., Fujita, H., Fukuda, Y., Noguchi, T., Doss, M., Kappas, A. and Sassa, S. (1992) Cloning and expression of the defective genes from a patient with delta-aminolevulinate dehydratase porphyria. J Clin Invest 89, 1431–7. 10.1172/JCI115732.

23. Dailey, H. A., Dailey, T. A., Gerdes, S., Jahn, D., Jahn, M., O’Brian, M. R. and Warren, M. J. (2017) Prokaryotic Heme Biosynthesis: Multiple Pathways to a Common Essential Product. Microbiology and Molecular Biology Reviews 81. 10.1128/mmbr.00048-16.

24. Layer, G., Reichelt, J., Jahn, D. and Heinz, D. W. (2010) Structure and function of enzymes in heme biosynthesis. Protein Science 19, 1137–1161. 10.1002/pro.405.

25. Hagiya, Y., Endo, Y., Yonemura, Y., Takahashi, K., Ishizuka, M., Abe, F., Tanaka, T., Okura, I., Nakajima, M., Ishikawa, T. and Ogura, S.ichiro (2012) Pivotal roles of peptide transporter PEPT1 and ATP-binding cassette (ABC) transporter ABCG2 in 5-aminolevulinic acid (ALA)-based photocytotoxicity of gastric cancer cells in vitro. Photodiagnosis Photodyn Ther 9, 204–214. 10.1016/j.pdpdt.2011.12.004.

26. Palasuberniam, P., Yang, X., Kraus, D., Jones, P., Myers, K. A. and Chen, B. (2015) ABCG2 transporter inhibitor restores the sensitivity of triple negative breast cancer cells to aminolevulinic acid-mediated photodynamic therapy. Sci Rep 5. 10.1038/srep13298.

27. Lubos, E., Loscalzo, J. and Handy, D. E. (2011) Glutathione peroxidase-1 in health and disease: From molecular mechanisms to therapeutic opportunities. Antioxid Redox Signal 15, 1957–1997. 10.1089/ars.2010.3586.

28. Pei, J., Pan, X., Wei, G. and Hua, Y. (2023) Research progress of glutathione peroxidase family (GPX) in redoxidation. Front Pharmacol 14. 10.3389/fphar.2023.1147414.

29. Arfin, S., Jha, N. K., Jha, S. K., Kesari, K. K., Ruokolainen, J., Roychoudhury, S., Rathi, B. and Kumar, D. (2021) Oxidative stress in cancer cell metabolism. Antioxidants 10. 10.3390/antiox10050642.

30. Huang, H. C., Mallidi, S., Liu, J., Chiang, C. Te Mai, Z., Goldschmidt, R., Ebrahim-Zadeh, N., Rizvi, I. and Hasan, T. (2016) Photodynamic therapy synergizes with irinotecan to overcome compensatory mechanisms and improve treatment outcomes in pancreatic cancer. Cancer Res 76, 1066–1077. 10.1158/0008-5472.CAN-15-0391.

31. Kessel, D. and Luo, Y. Photodynamic therapy: A mitochondrial inducer of apoptosis. 10.8.98.

32. Oleinick, N. L., Morris, R. L. and Belichenko, I. (2002) The role of apoptosis in response to photodynamic therapy: What, where, why, and how. Photochemical and Photobiological Sciences 1, 1–21. 10.1039/b108586g.

33. Kessel, D. and Luo, Y. (1998) Mitochondrial photodamage and PDT-induced apoptosis. J Photochem Photobiol B 42, 89–95. 10.1016/s1011-1344(97)00127-9.

34. Anand, S., Wilson, C., Hasan, T. and Maytin, E. V. (2011) Vitamin D3 enhances the apoptotic response of epithelial tumors to aminolevulinate-based photodynamic therapy. Cancer Res 71, 6040–6050. 10.1158/0008-5472.CAN-11-0805.

35. Rizvi, I., Celli, J. P., Evans, C. L., Abu-Yousif, A. O., Muzikansky, A., Pogue, B. W., Finkelstein, D. and Hasan, T. (2010) Synergistic enhancement of carboplatin efficacy with photodynamic therapy in a three-dimensional model for micrometastatic ovarian cancer. Cancer Res 70, 9319–9328. 10.1158/0008-5472.CAN-10-1783.

36. Struth, E., Labaf, M., Karimnia, V., Liu, Y., Cramer, G., Dahl, J. B., Slack, F. J., Zarringhalam, K., & Celli, J. P. (2024). Drug resistant pancreatic cancer cells exhibit altered biophysical interactions with stromal fibroblasts in imaging studies of 3D co-culture models. bioRxiv. 10.1101/2024.07.14.602133

